# Discovery and optimization of piperazine-1-thiourea-based human phosphoglycerate dehydrogenase inhibitors

**DOI:** 10.1101/250340

**Authors:** Jason M. Rohde, Kyle R. Brimacombe, Li Liu, Michael E. Pacold, Adam Yasgar, Dorian M. Cheff, Tobie D. Lee, Ganesha Rai, Bolormaa Baljinnyam, Zhuyin Li, Anton Simeonov, Matthew D. Hall, Min Shen, David M. Sabatini, Matthew B. Boxer

## Abstract

Proliferating cells, including cancer cells, obtain serine both exogenously and via the metabolism of glucose. By catalyzing the first, rate-limiting step in the synthesis of serine from glucose, phosphoglycerate dehydrogenase (PHGDH) controls flux through the biosynthetic pathway for this important amino acid and represents a putative target in oncology. To discover inhibitors of PHGDH, a coupled biochemical assay was developed and optimized to enable high-throughput screening for inhibitors of human PHGDH. Feedback inhibition was minimized by coupling PHGDH activity to two downstream enzymes (PSAT1 and PSPH), providing a significant improvement in enzymatic turnover. Further coupling of NADH to a diaphorase/resazurin system enabled a red-shifted detection readout, minimizing interference due to compound autofluorescence. With this protocol, over 400,000 small molecules were screened for PHGDH inhibition, and following hit validation and triage work, a piperazine-1-thiourea was identified. Following rounds of medicinal chemistry and SAR exploration, two probes (NCT-502 and NCT-503) were identified. These molecules demonstrated improved target activity and encouraging ADME properties, enabling both *in vitro* and *in vivo* assessment of the biological importance of PHGDH, and its role in the fate of serine in PHGDH-dependent cancer cells.

## Introduction

Serine is an important biochemical building block, being incorporated wholly into proteins as well as into DNA and RNA bases as one-carbon units or as serine-derived glycine. Given this crucial role, proliferating cells obtain serine exogenously or derive it from glucose. By catalyzing the first, rate-limiting step in the synthesis of serine from glucose, phosphoglycerate dehydrogenase (PHGDH) controls flux through this biosynthetic pathway. A series of PHGDH-overexpressing breast cancer cell lines display a dependence on this pathway as genetic suppression of PHGDH was shown to be toxic even in the presence of exogenous serine.^1–2^

Given PHGDH’s potential as a drug target in certain cancers, a number of PHGDH inhibitors have been reported.^3–11^ AstraZeneca used a fragment-based lead discovery approach followed by medicinal chemistry optimization to develop an indole derivative (Figure 1) with submicromolar K_D_.^4–5^ Cantley and colleagues used a biochemical high-throughput screen to identify a PHGDH inhibitor (CBR-5884, Figure 1) with an IC_50_ of ∼33 ìM.^6^ A patent filed by Raze Therapeutics disclosed a chemotype (representative example shown in Fig 1), but no detailed information on inhibitory activity was shown.^9^

**Figure 1.**
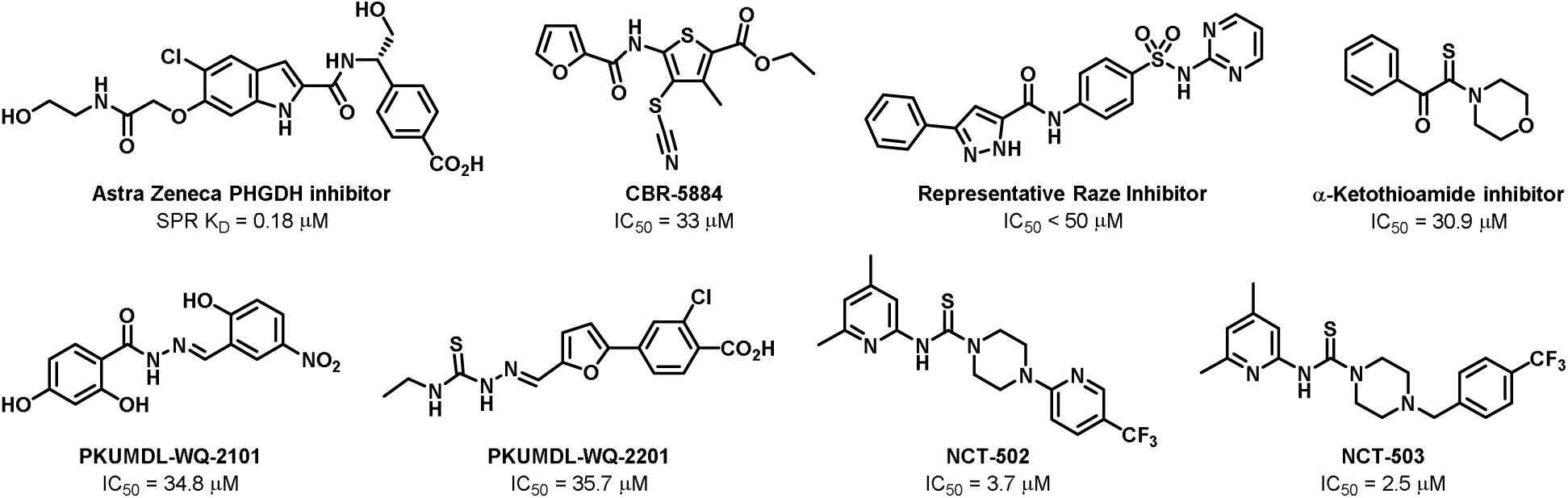
Reported PHGDH inhibitors

We have previously reported inhibitors of PHGDH (NCT-502, NCT 503, Figure 1) with low-micromolar biochemical inhibition that demonstrated inhibition of PHGDH-dependent cancer cell growth, and reduced glucose-derived serine production in cells.^8^ Activity was also demonstrated in orthotopic xenograft tumors.

Here, we describe the development and optimization of a high-throughput amenable PHGDH biochemical activity assay to enable screening in 1536-well format. This assay was utilized to conduct a quantitative high-throughput screen (qHTS) to uncover small molecule inhibitors of PHGDH for further development. In addition, we describe the optimization of a thiourea chemotype into a set of *in vitro* and *in vivo* probes to study PHGDH biology.

## Results

The optimized PHGDH assay used for the HTS utilized a coupled enzyme system to provide robust recombinant PHGDH activity and to shift detection of dehydrogenase activity from a classic NADH-based blue fluorescent readout to a red-shifted resorufin-based readout. This coupled biochemical assay system includes two human enzymes – phosphoserine transaminase (PSAT1) and phosphoserine phosphatase (PSPH) – which naturally participate in the serine biosynthetic pathway immediately downstream of PHGDH, along with a bacterial enzyme, diaphorase from *Clostridium kluyveri*, to couple NADH levels to the resazurin/resorufin fluorescent dye system (Figure 2a).^12^

**Figure 2.**
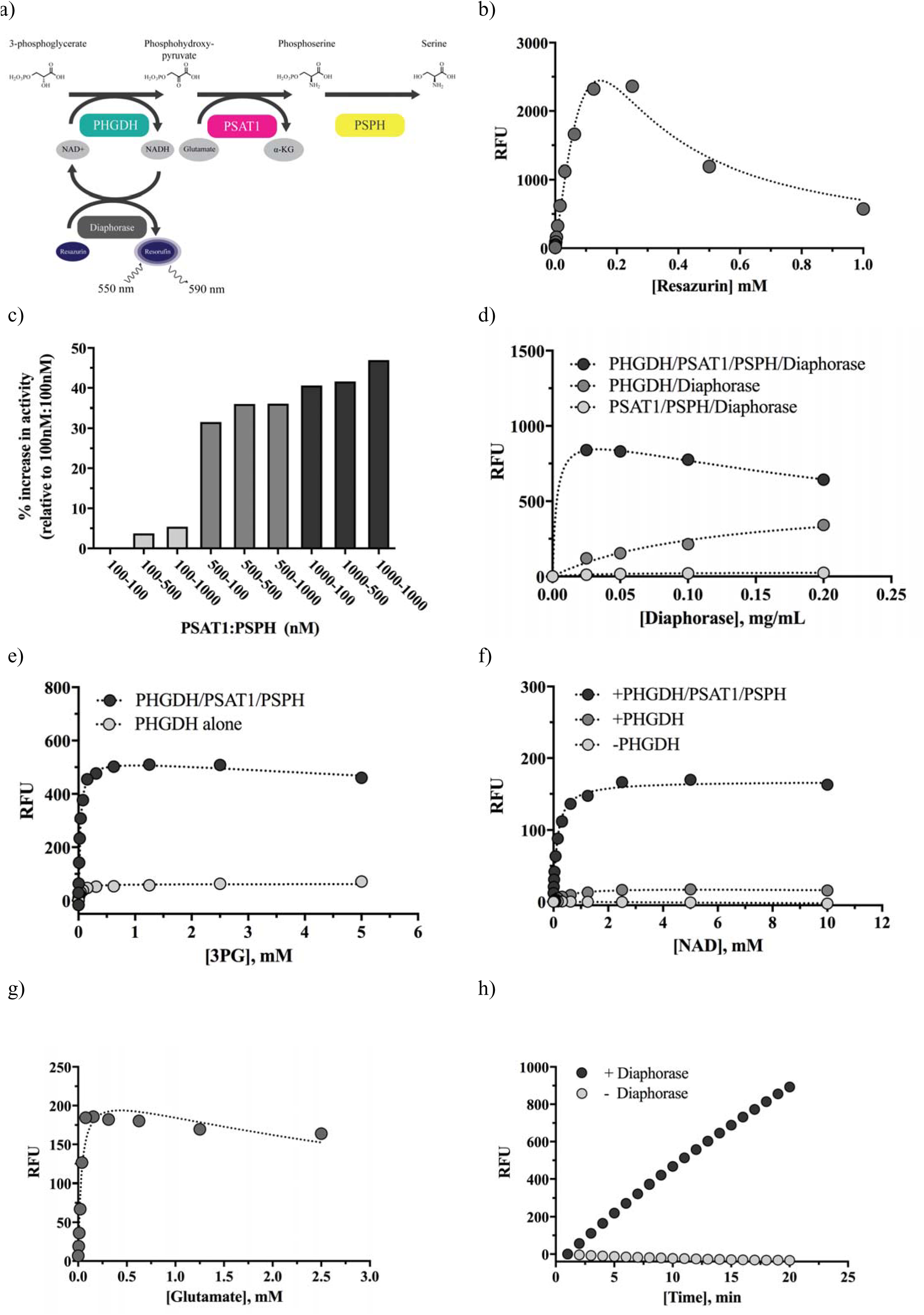
PHGDH assay development. a) Scheme of PHGDH enzymatic reaction coupled to diaphorase for fluorescence detection. b) The diaphorase assay was run using a resazurin titration to determine the substrate’s Km (0.05 mM), and feedback inhibition was observed at substrate concentrations above 0.25 mM. c) PHGDH activity (10 nM) was measured utilizing varying stoichiometries of PSAT1 and PSPH to relative contributions to PHGDH assay turnover. Conditions for the coupled PHGDH assay were determined by testing titrations of d) diaphorase, e) 3PG, f) NAD^+^ and g) glutamate to identify ideal assay parameters (i.e. Kms for the substrates 3PG and NAD^+^, and non-limiting concentrations for the coupling reagents diaphorase and glutamate). h) A time course demonstrating robust, near-linear kinetics is shown for the fully-coupled assay in the presence and absence of diaphorase.

### Assay Development & Optimization

The coupling efforts detailed here were undertaken after initial adaptation of a PHGDH assay containing solely the dehydrogenase and its substrates, which quantified NADH production by directly measuring NADH fluorescence, yielded low enzymatic turnover and required a lengthy reaction period to achieve significant signal (data not shown). Furthermore, given the prevalence of blue autofluorescence among small molecules in screening collections, we anticipated a need to shift the primary assay away from the blue NADH-based fluorescent readout.^13^

To address these issues, we coupled NADH production to diaphorase, which has been utilized in other dehydrogenase and NAD^+^/NADH systems to introduce a separate, red-shifted detection dye, resazurin.^14^ Diaphorase is an oxidoreductase enzyme that can utilize NADH or NADPH to catalyze the reduction of resazurin to fluorescent resorufin. Diaphorase-coupling provides a twofold benefit in the case of PHGDH by 1) generating resorufin, a robust detection dye with HTS-amenable red-shifted fluorescence, with 1:1 stoichiometry to NADH, and by 2) regenerating NAD^+^ and preventing accumulation of NADH by the primary PHGDH reaction (which can slow enzyme kinetics). We tested *Clostridium kluyveri* diaphorase for this purpose by determining optimal resazurin and diaphorase concentrations using PHGDH assay buffer and substrate conditions. Titration of resazurin using 0.5 mM NADH and 0.015 mg/mL diaphorase yielded a K_m_ of 50 μM for resazurin, and determined that maximal production of resorufin signal occurred at 250 μM substrate; concentrations above this threshold were found to demonstrate feedback inhibition with reduced signal (Figure 2b). For coupling purposes, resazurin was included at 0.1 mM, which was high enough to achieve a non-limiting concentration relative to NAD^+^/NADH in the enzyme system, yet low enough to avoid any potential feedback inhibition.

PHGDH has been shown to be susceptible to feedback inhibition^15^ by its product phosphohydroxypyruvate (p-Pyr), so the assay system was further coupled to two downstream enzymes, PSAT1 and PSPH, to minimize p-Pyr accumulation and help ‘drive’ the reaction forward. Optimal stoichiometry of PHGDH, PSAT1 and PSPH were determined by titrating the downstream enzymes PSAT1 and PSPH in the presence of 10 nM PHGDH and non-limiting substrate concentrations (1 mM NAD^+^, 2 mM 3-phosphoglycerate [3PG], 0.625 mM glutamate), using NADH fluorescence as a direct readout of biochemical reaction progression. Inclusion of PSAT1 and PSPH led to increased PHGDH turnover and greater assay signal at all tested concentrations and stoichiometries, though PSAT1 concentrations were found to most strongly influence overall assay signal (Figure 2c). Inclusion of 500 nM PSAT1 and 500 nM PSPH yielded a 36% increase in signal over the lowest concentrations tested (100 nM each); though higher PSAT1 and PSPH concentrations did provide further increases in signal. Therefore, 500 nM concentrations of both enzymes were chosen to balance assay signal with protein requirements.

Next, we performed a diaphorase titration in the presence of either PHGDH alone, PHGDH with PSAT1 and PSPH (positive control), or PSAT1 and PSPH alone (no PHGDH, negative control). The fully-coupled PHGDH/PSAT1/PSPH/diaphorase assay conditions yielded significantly greater assay signal and turnover than other conditions, with 2-7-fold increased turnover compared to PHGDH/diaphorase in the absence of PSAT1 and PSPH (Figure 2d), confirming that the coupled assay increases PHGDH enzymatic activity. Though maximal assay signal was seen at 0.025 mg/mL diaphorase in the fully-coupled conditions, diaphorase was included in subsequent screening at a higher concentration (0.1 mg/mL) to buffer against any significant false positive readings in the event of direct inhibition of diaphorase itself.

With coupling conditions determined, the assay system substrates 3PG, NAD^+^ (PHGDH) and glutamate (PSAT1) were individually titrated to determine K_m_ values and saturating concentrations. K_m_ values for 3PG and NAD^+^ were determined to be 50 μM and 150 μM, respectively, and these concentrations were adopted into the final assay conditions to facilitate detection of small molecule competitors of either substrate (Figures 2e, f). Glutamate was found to be non-limiting above 310 μM, so 625 μM glutamate was utilized in the final assay to ensure saturation for the PSAT1/PSPH coupling reaction (Figure 2g). The final, fully-coupled PHGDH assay demonstrated robust, linear turnover over 20 minutes, suggesting it was amenable to screening (Figure 2h). These conditions represent the final coupled PHGDH assay utilized to conduct all subsequent screening and validation work (Table 1).

**Table 1.** Primary PHGDH coupled assay protocol

| Step |  | Value | Description |
| --- | --- | --- | --- |
| 1 | Reagent Addition | 3 μL | Substrate mix in assay buffer |
| 2 | Compound Addition | 23 nL | Dilution series (4-11 point series in DMSO) |
| 3 | Reagent Addition | 1 μL | PHGDH enzyme mix in assay buffer |
| 4 | Detection | Fluorescence | ViewLux initial read, T0 min |
| 5 | Incubation | 20 min | Room temperature incubation |
| 6 | Detection | Fluorescence | ViewLux final read, T20 min |
| Notes |  |  |  |
| 1 | Substrate buffer: 50 mM triethanolamine, 10 mM MgCl<sub>2</sub>, 0.05% BSA, 0.01% Tween20, 10 μM EDTA, 0.625 mM glutamate, 0.05 mM 3-phosphoglycerate, 0.1 mM resazurin, 500 nM PSAT1, 500 nM PSPH and 0.1 mg/mL diaphorase | | |
| 2 | Pintool transfer |  |  |
| 3 | PHGDH enzyme buffer: 50 mM TEA, 10 mM MgCl<sub>2</sub>, 0.05% BSA, 0.01% Tween20, 0.15 mM NAD<sup>+</sup>, 10 nM PHGDH |  |  |
| 4 | ViewLux fluorescence read (ex525/em598, 5 sec exposure, 5000 excitation energy) |  |  |
| 5 | Room temperature |  |  |
| 6 | ViewLux fluorescence read (ex525/em598, 5 sec exposure, 5000 excitation energy) |  |  |

### Pilot Screening

To compare the performance of the optimized assay to the original uncoupled version, preliminary screening was conducted using the LOPAC^®1280^ library. In this pilot screen, the uncoupled NADH assay resulted in interference from significant compound fluorescence, which was of a considerably greater magnitude than the NADH fluorescence resulting from PHGDH turnover and NADH production (Figure 3a).

**Figure 3.**
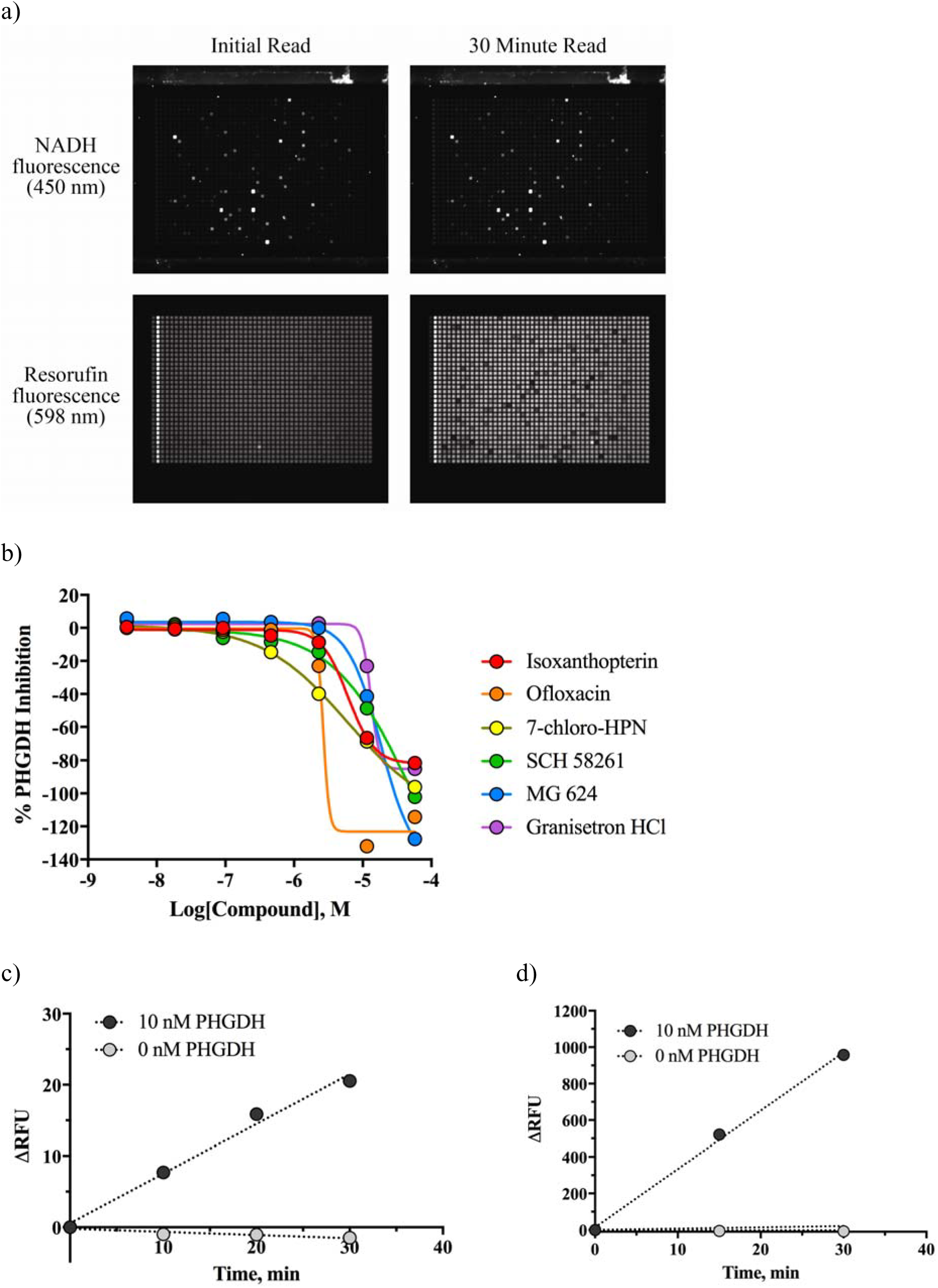
PHGDH preliminary screening and optimization. a) Plate images are shown for T=0 (left plates) and T=30 min (right plates) reads from two PHGDH screens against the LOPAC library (11.5 μM). The two top plates demonstrate the autofluorescence of LOPAC library compounds in the PHGDH assay utilizing NADH-based detection, while the bottom two plates show well signal from the PHGDH assay using resazurin-based detection. Very few autofluorescent compounds can be observed in the resazurin wavelength, providing a clearer landscape for observing potential PHGDH inhibition. b) Examples are shown of false-positive hits identified from LOPAC in the PHGDH assay utilizing NADH-based detection. All corresponding compounds were completely inactive in the diaphorase/resazurin-based PHGDH assay, demonstrating an advantage of migrating to the 598 nm detection realm. Assay signal windows observed during LOPAC screens using c) NADH-linked or d) resorufin-based readouts.

Using Δ30 min data at 57 μM compound, where inhibition ≥50% was defined as active, the NADH-fluorescence assay yielded 103 hits out of 1280 compounds (hit rate of 8%). In contrast, the PSAT1/PSPH/diaphorase-coupled assay showed nearly 3-fold fewer autofluorescent compound artifacts, (37 out of 1280 compounds, hit rate of 3%) and at a comparatively lower magnitude relative to resorufin signal. During these screens, plates were measured every 10-15 min for a total of 30 min, and 32 positive control (10 nM PHGDH enzyme) and negative control (no PHGDH enzyme) wells were included on each 1536-well plate to monitor assay performance. The uncoupled assay (direct NADH detection) yielded a ΔRFU of only 20 after 30 minutes (Figure 3c), while the diaphorase assay demonstrated a ΔRFU of nearly 1000, representing a 50-fold increase in signal over the same time period (Figure 3d). The coupled assay also demonstrated superior performance to the original assay, for both signal-to-background (5.1 vs 3.7) and Z’ (0.85 vs 0.80).

### Primary Quantitative High-throughput Screening

The coupled PHGDH assay was utilized to screen our in-house collection of more than 400,000 small molecules. In this qHTS screen, the inhibition associated with each well was computed from the endpoint and normalized against control wells. The percent inhibition at each of the concentrations of inhibitor tested was fit to a sigmoidal dose-response curve (Hill equation) using in-house-developed software (https://tripod.nih.gov/curvefit) to determine the compound IC_50_ values. Across >500 1536-well plates, assay performance was strong with Z’ generally >0.7 and S:B >20 (Figure 4a). Analysis of these dose-response curves resulted in 1342 high-quality actives that showed strong inhibition with curve classes 1-3, efficacy over 50% and IC_50_ ≤ 20 μM (Figure 4b). These hit compounds were then thoroughly evaluated for both promiscuity and synthetic tractability to eliminate compounds that did not offer good starting points for lead optimization. A total of 239 prioritized inhibitors were reacquired and screened in 11-point dose-response in the primary screening assay to confirm their activity.

**Figure 4.**
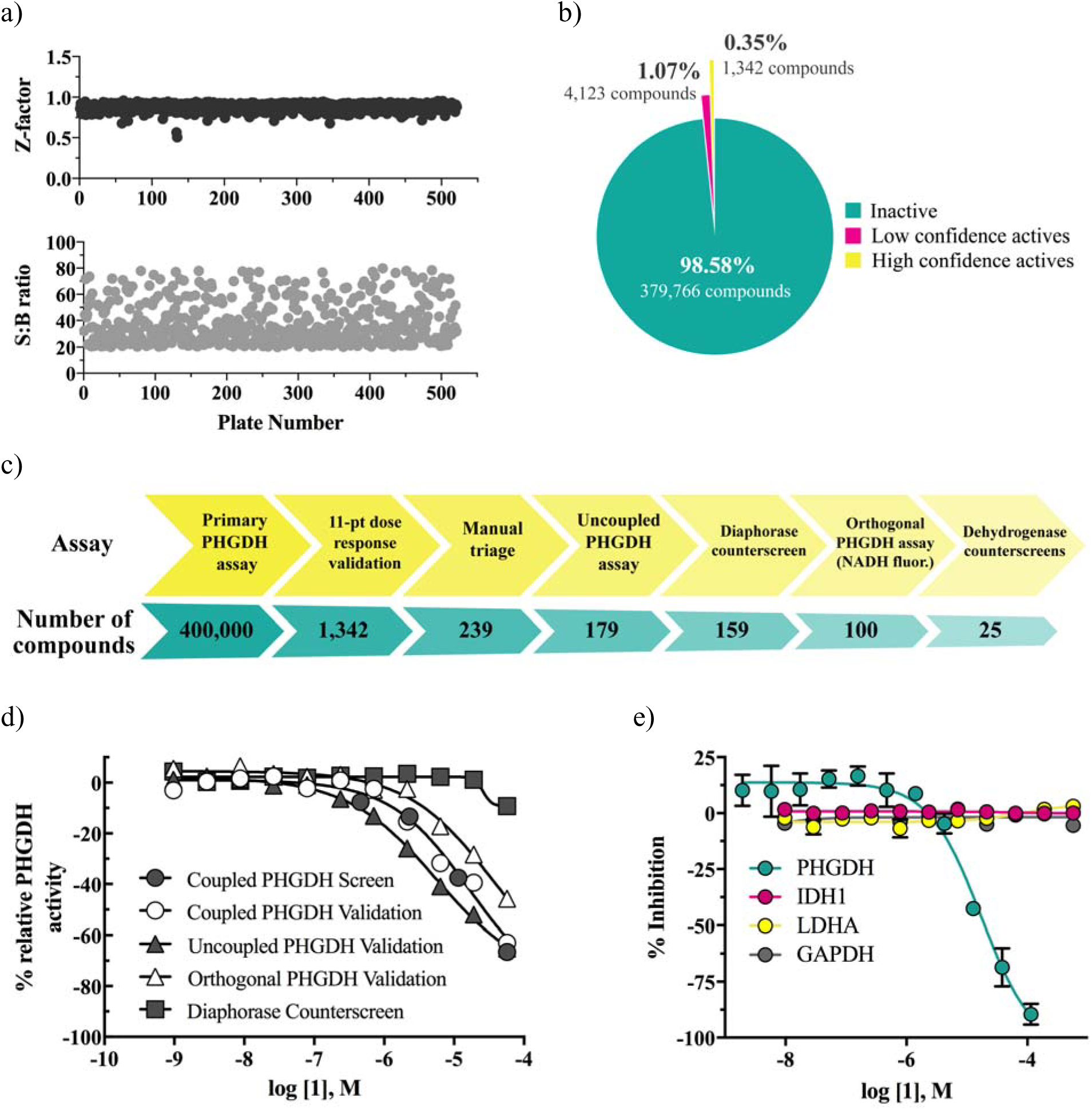
PHGDH high-throughput screen results. a) Assay performance statistics (Z-factor, top, and signal-to-background ratio, bottom) are shown for each plate across the entire PHGDH qHTS campaign. b) A breakdown of the number and quality of qHTS hits is shown. High confidence actives are those demonstrating curve classes of −1.1, −1.2 and −2.1, while low confidence actives demonstrated curve classes of −1.3, −1.4, −2.2 or −3. c) The PHGDH assay hit triage funnel, showing the assays used to winnow the initial collection of screening hits, along with the number of compounds remaining after each stage. d) Dose-response data showing activity of screening hit (**1**) in the assay hit triage funnel. Data is shown for **1** in the primary coupled screen and cherry-pick validation (PHGDH, PSAT1, PSPH and diaphorase), the uncoupled secondary assay (PHGDH and diaphorase), the orthogonal secondary assay (PHGDH, PSAT1 and PSPH), and the diaphorase-only counterscreen assay. e) Dose-response data demonstrating selectivity of the screening hit **1** against a panel of dehydrogenase assays (PHGDH, IDH1, LDHA and GAPDH).

### Hit Validation and Triage

A number of secondary assays and counterscreens were selected and utilized to refine and triage initial screening hits (assay hit triage funnel shown in Figure 4c). This triage panel included 1) validation in the primary assay using expanded 11-point dose-response, 2) triage of PAINS^16–17^ or other chemotypes/scaffolds not amenable to medicinal chemistry optimization, 3) orthogonal validation in the original uncoupled PHGDH assay (PHGDH alone, using NADH fluorescence), 4) counterscreens against diaphorase alone, 5) orthogonal screening against the coupled PHGDH/PSAT1/PSPH reaction using NADH fluorescence (without diaphorase), and 6) counterscreens against unrelated dehydrogenases (isocitrate dehydrogenase [IDH1], lactate dehydrogenase A [LDHA] and glyceraldehyde 3-phosphate dehydrogenase [GAPDH]).

The activity of **1** was subsequently characterized in dose-response using the outlined assay triage hit funnel (Figure 4d). The primary screen and secondary cherry-pick validation activity of **1** was found to be in close agreement (14.1 μM vs 15.3 μM). In the uncoupled PHGDH assay, which utilized diaphorase and resazurin for detection without PSAT1 and PSPH, **1** displayed an increase in potency (5.4 μM), suggesting that its activity was independent of the presence/activity of the downstream enzymes. Similarly, **1** retained activity in the orthogonal PHGDH assay (17.2 μM), which included PSAT1 and PSPH but utilized NADH fluorescence as a direct measure of turnover, demonstrating that its activity was not dependent on the diaphorase/resazurin coupling system. This observation was further confirmed in the diaphorase counterscreen assay, which showed no appreciable inhibition of diaphorase (<9.3% inhibition at the top concentration of 57.5 μM). Taken together, the assay panel suggested that **1**’s activity was directly dependent on PHGDH, rather than any of the coupling enzymes.

Select hits were advanced for selectivity testing, which included a small panel of dehydrogenases (namely IDH1, LDHA and GAPDH) to exclude pan-dehydrogenase inhibitors. A number of hit compounds were found to demonstrate strong selectivity toward PHGDH over other dehydrogenases, with little comparative off-target activity. Notably, the screening hit **1** displayed complete selectivity toward PHGDH, with no discernable off-target dehydrogenase inhibition (Figure 4e).

### Medicinal Chemistry & SAR Exploration

Based on this panel of assays, a series of 2-pyridinyl-*N*-(4-aryl)piperazine-1-carbothioamides emerged as the top, validated chemotype, as low micromolar inhibitors of PHGDH. This chemotype was explored earlier during another NCGC structure-activity campaign to develop inhibitors of bacterial phosphopantetheinyl transferase (PPTase)^18^. Utilizing compounds from this previous medicinal chemistry effort, we were able to rapidly establish preliminary structure-activity relationships (SAR) beyond hit **1** (Table 2).

**Table 2.** Initial SAR focusing on the thiourea group of hit **1**

| Cmpd | R | IC<sub>50</sub>(μM)<sup>a</sup> | % inhibition |
| --- | --- | --- | --- |
| 1 |  | 15.3 | 92 |
| 2 |  | ND | 38 (max) |
| 3 |  | ND | 16 (max) |
| 4 |  | ND | 2 (max) |
| 5 |  | ND | 37 (max) |
| 6 |  | ND | 9 (max) |
| 7 |  | 21 | 45 |

<sup>a</sup>IC<sub>50</sub> values were determined utilizing the diaphorase and resazurin-coupled PHGDH assay. ND = not determined; for compounds with % inhibition at 57 μM weaker than 40%, IC<sub>50</sub> was ND.

We began our PHGDH SAR investigations with a focus on the necessity of the thiourea group (Table 2). This feature of the pharmacophore proved essential to activity as the urea (**3**), guanidine (**4**), and thioamide (**5**) replacements led to significant loss of PHGDH activity. Additionally, other minor modifications to this region of the molecule were not tolerated such as *N*-methylation (**2**), shortening of the linkage from the thiocarbonyl to the pyridine (**6**), or lengthening the linkage (**7**).

Subsequently, we shifted our SAR analysis to the aminopyridine portion of the chemotype (Table 3). The pyridine in this region was required to maintain target inhibition as phenyl variant **8** lost all potency. The positioning of the pyridine nitrogen also proved to be optimal at the two-position as the 3− and 4-pyridyl analogs **10** and **11** lost both potency and efficacy in comparison to compound **9**. A second nitrogen proximal to the thiourea nitrogen in the form of a pyrimidine (**12**) led similarly to diminished PHGDH activity as did substitution of the pyridine with electron-withdrawing groups (analogs **13** and **14**). The addition of a second methyl group resulted in an improvement in potency, whereas expansion of the pyridine ring system in the form of a quinoline (analogs **16** and **17**) showed small perturbations in target activity.

**Table 3.** Brief SAR of the aminopyridine portion of hit **1**

| Cmpd | R | IC<sub>50</sub>(μM)<sup>a</sup> | % inhibition | Cmpd | R | IC<sub>50</sub>(μM)<sup>a</sup> | % inhibition |
| --- | --- | --- | --- | --- | --- | --- | --- |
| 1 |  | 15.3 | 92 | 13 |  | 6.4 | 61 |
| 8 |  | NA | 20 (max) | 14 |  | 13 | 73 |
| 9 |  | 6.8 | 93 | 15 |  | 6.3 | 92 |
| 10 |  | 8.9 | 67 | 16 |  | 10 | 90 |
| 11 |  | 14 | 55 | 17 |  | 21 | 119 |
| 12 |  | 37 | 111 |  |  |  |  |

<sup>a</sup>IC<sub>50</sub> values were determined utilizing the diaphorase and resazurin-coupled PHGDH assay. ND = not determined; for compounds with % inhibition at 57 μM weaker than 40%, IC<sub>50</sub> was ND.

A key feature of our approach to the optimization of this chemotype involved the evaluation of structure-property relationships (SPR) in parallel to all of our SAR work. We performed a set of *in vitro* absorption, distribution, metabolism and excretion (ADME) assays on all compounds including aqueous kinetic solubility, rat liver microsomal stability (single point), and parallel artificial membrane permeability assay (PAMPA). At this stage of our investigations, we leveraged improved SPR observations as compound **15** displayed improved microsomal stability (rat liver microsomes t_1/2_ = >30 min) compared to analog **9** (t_1/2_ = 16 min). Thus, dimethylpyridine was carried forward for subsequent SAR studies on the remaining regions of the chemotype. Furthermore, although only a small sample of analogs probing this section of the hit molecule **1** are displayed here, a broader exploration carried out later in the SAR campaign confirmed, in the same way, that this chemotype would not accommodate alterations to the pyridyl ring system.

Having established that the 2-pyridine and the thiourea are important features of the pharmacophore, we shifted our attention to the arene affixed to the piperazine in order to assess the flexibility therein (Table 4). Movement of the trifluoromethyl group around the arene proved to be well-tolerated with the best location being *para* to the piperazine (**19**). Unsubstituted pyridines lost most of their PHGDH activity regardless of their positioning (analogs **20**-**22**). Given that analog **22** maintains many structural features of the hit **1** as well as subsequent molecules, it became incorporated into *in vitro* studies as an inactive control. Further explorations with electron-donating methoxy groups in this region led to modest (**24**, **25**) to significant (**23**) losses in potency depending on the positioning. Lastly, in an attempt to boost solubility, we combined the trifluoromethyl substituent with the pyridine in the most tolerant position to generate analogs **26** and **27** (NCT-502). While NCT-502 (**27**) benefitted from a small improvement in PHGDH potency, it did not benefit from an increase in kinetic aqueous solubility (1.23 μM at pH 7.4) as we expected. Despite its relatively poor aqueous solubility, NCT-502 (**27**) proved to be a valuable tool molecule which was used to begin understanding the *in vitro* biology of human PHGDH.

**Table 4.** Further SAR of the *N*-aryl piperazine of compound **15** and discovery of NCT-502 (**27**)

| Cmpd | R | IC<sub>50</sub>(μM)<sup>a</sup> | % inhibition |
| --- | --- | --- | --- |
| 15 |  | 6.3 | 92 |
| 18 |  | 8.7 | 101 |
| 19 |  | 4.3 | 98 |
| 20 |  | 37 | 73 |
| 21 |  | 50 | 68 |
| 22 |  | NA | 33 (max) |
| 23 |  | 24 | 109 |
| 24 |  | 14 | 101 |
| 25 |  | 16 | 99 |
| 26 |  | 11 | 100 |
| 27<br>(NCT-502) |  | 3.7 | 96 |

<sup>a</sup>IC<sub>50</sub> values were determined utilizing the diaphorase and resazurin-coupled PHGDH assay. ND = not determined; for compounds with % inhibition at 57 μM weaker than 40%, IC<sub>50</sub> was ND.

**Table 5.** Core SAR of compound **15** and discovery of NCT-503 (**31**)

| Cmpd |  | R | IC<sub>50</sub>(μM)<sup>a</sup> | % inhibition |
| --- | --- | --- | --- | --- |
| 15 |  | *m*-CF<sub>3</sub>Ph | 6.3 | 92 |
| 19 |  | *p*-CF<sub>3</sub>Ph | 4.3 | 98 |
| 28 |  | *m*-CF<sub>3</sub>benzoyl | 22 | 101 |
| 29 |  | *p*-CF<sub>3</sub>benzoyl | 19 | 85 |
| 30 |  | *m*-CF<sub>3</sub>benzyl | 5.3 | 102 |
| 31<br>(NCT-503) |  | *p*-CF<sub>3</sub>benzyl | 2.5 | 96 |
| 32 |  | *m*-CF<sub>3</sub>Ph | 7.2 | 84 |
| 33 |  | *p*-CF<sub>3</sub>Ph | 5.7 | 85 |
| 34 |  | *m*-CF<sub>3</sub>benzoyl | 10 | 91 |
| 35 |  | *p*-CF<sub>3</sub>benzoyl | 8.7 | 88 |
| 36 |  | *p*-CF<sub>3</sub>benzyl | 12 | 89 |
| 37 |  | *m*-CF<sub>3</sub>Ph | 6.5 | 90 |
| 38 |  | *p*-CF<sub>3</sub>Ph | 7.7 | 97 |
| 39 |  | *m*-CF<sub>3</sub>benzoyl | 11 | 94 |
| 40 |  | *p*-CF<sub>3</sub>benzoyl | 10 | 90 |
| 41 |  | *p*-CF<sub>3</sub>benzyl | 15 | 93 |

<sup>a</sup>IC<sub>50</sub> values were determined utilizing the diaphorase and resazurin-coupled PHGDH assay. ND = not determined; for compounds with % inhibition at 57 μM weaker than 40%, IC<sub>50</sub> was ND.

With insufficient improvements in target potency and *in vitro* ADME properties through early SAR and SPR studies, we considered more significant modifications to the piperazine core in conjunction with changes to the group bridging to the trifluoromethyl phenyl ring. Along these lines, we installed a one-carbon spacer (both methylene and carbonyl) from the piperazine core to the arene segment. Additionally, we excised the piperazine nitrogen distal to the thiourea to a pair of amino piperidines (both 3− and 4-substituted). Methods to access many of these differentially substituted cores were reported previously.^8, 18^ The majority of the piperazine analogs maintained much of their activity (**15**, **19**, **30**, **31**) with significant drop-offs being observed only with amides **28** and **29**. The optimal 3-amino piperidine analog was that of the *p*trifluoromethyl aniline **33**. The other changes to this core (**32**, **34-36**) resulted in modest decreases in activity. A similar trend was observed for the 4-amino piperidine core, with analogs **37** and **38** proving to be the best among the set, and analogs **39-41** showing losses in activity. With regard to the three linkers, benzoyl led to a moderate drop in potency across all three cores. Interestingly, the benzyl linkage proved optimal with the original piperazine core (analogs **30** and **31** [NCT-503]) as more significant drops in potency were observed with the piperidine ring system (compounds **36** and **41**). Lastly, the piperazine core was chosen for further SAR evaluation as the solubility of top analog, **31** [NCT-503] (aqueous solubility = 25.3 μM at pH 7.4), was much greater than that of the top analogs from each of the amino piperidine sets (analogs **33** and **37** displayed aqueous kinetic solubilities below the limits of quantitation).

To further probe the ethylene diamine feature of the core, a subset of ring-opened analogs were evaluated. The increased flexibility resulted in a loss of activity with the dimethylated analog **44** maintaining moderate potency (Table 6). Conversely, rigid variations of the core, such as the fused *bis*-pyrrolidine analog **45** and 1,4-diazepane **46**, were also less active than NCT-503 (**31**). Returning to the piperazine and the methylene linkage to the trifluoromethyl arene as above, we wanted to investigate whether substitution (**47**) or extension (**48**) would improve potency. Unfortunately, both changes resulted in diminished activity. Furthermore, similar to much of our previous SAR, piperazine substitution with both hydrophobic groups (**49-52**, **55**) as well as hydrophilic groups (**53**, **54**) proved deleterious, with analogs **49** and **53** experiencing only small decreases in PHGDH inhibition.

**Table 6.** Piperazine SAR of compound **31** (NCT-503)

| Cmpd | | R | IC<sub>50</sub>(μM)<sup>a</sup> | % inhib. | Cmpd | | IC<sub>50</sub>(μM)<sup>a</sup> | % inhib. |
| --- | --- | --- | --- | --- | --- | --- | --- | --- |
| 42 |  | R = *p*-CF<sub>3</sub>benzyl | 9.7 | 41 | 49 |  | 8.3 | 99 |
| 43 |  | R = *p*-CF<sub>3</sub>benzyl | 21 | 66 | 50 |  | 12 | 94 |
| 44 |  | R = *p*-CF<sub>3</sub>benzyl | 7.3 | 84 | 51 |  | 9.9 | 86 |
| 45 |  | R = *p*-CF<sub>3</sub>benzyl | 16 | 93 | 52 |  | NA | 16 (max) |
| 46 |  | R = *p*-CF<sub>3</sub>benzyl | 7.3 | 82 | 53 |  | 5.8 | 89 |
| 47 |  | R = | 8.0 | 90 | 54 |  | 33 | 93 |
| 48 |  | R = CH<sub>2</sub>CH<sub>2</sub>(*p*-CF<sub>3</sub>phenyl) | 23 | 87 | 55 |  | 31 | 111 |

<sup>a</sup>IC<sub>50</sub> values were determined utilizing the diaphorase and resazurin-coupled PHGDH assay. ND = not determined; for compounds with % inhibition at 57 μM weaker than 40%, IC<sub>50</sub> was ND.

With the final phase of our SAR efforts focusing on a systematic exploration of the piperazine aryl group, we hoped a thorough R group scan may provide a breakthrough in terms of target potency. The general methodology we employed for this stage of the SAR campaign is highlighted in the context of the synthesis of NCT-503 (**31**) (Scheme 1).^1^ Selective *N-*benzylation of Boc-protected piperazine, protecting group removal, and subsequent one-pot coupling to *in situ*-generated *N*-(4,6-dimethylpyridin-2-yl)-1H-imidazole-1-carbothioamide proved direct and efficient. Thus, we scanned the positioning of many common aromatic substituents about this ring (Me, OMe, F, Cl, CF_3_) without improvements in PHGDH activity (Table 7). Testing of this large set of analogs focused on this aryl region of the chemotype proved unable to improve potency.

**Table 7.** Systematic SAR of benzyl piperazine portion of the pharmacophore

| Cmpd | Ar | IC<sub>50</sub>(μM)<sup>a</sup> | % inhib. |
| --- | --- | --- | --- |
| **56** | phenyl | **40** | **78** |
| **57** | 2-pyridyl | **50** | **51** |
| **58** | 3-pyridyl | **NA** | **34 (max)** |
| **59** | 4-pyridyl | **59** | **67** |
| **60** | *o*-tolyl | **9.2** | **85** |
| **61** | *m*-tolyl | **20** | **92** |
| **62** | *p*-tolyl | **17** | **93** |
| **63** | *o*-anisoyl | **NA** | **21 (max)** |
| **64** | *m*-anisoyl | **30** | **94** |
| **65** | *p*-anisoyl | **26** | **87** |
| **66** | *o*-fluorophenyl | **41** | **56** |
| **67** | *m*-fluorophenyl | **39** | **88** |
| **68** | *p*-fluorophenyl | **38** | **86** |
| **69** | *o*-chlorophenyl | **20** | **106** |
| **70** | *m*-chlorophenyl | **11** | **90** |
| **71** | *p*-chlorophenyl | **15** | **97** |
| **72** | *o*-trifluoromethylphenyl | **6.3** | **92** |
| **30** | *m*-trifluoromethylphenyl | **5.3** | **102** |
| **31**<br>(NCT-503) | *p*-trifluoromethylphenyl | **2.5** | **96** |
| **73** | *p*-bromophenyl | **17** | **92** |
| **74** | *p*-trifluoromethoxyphenyl | **11** | **96** |

<sup>a</sup>IC<sub>50</sub> values were determined utilizing the diaphorase and resazurin-coupled PHGDH assay. ND = not determined; for compounds with % inhibition at 57 μM weaker than 40%, IC<sub>50</sub> was ND.

## Discussion

The assay design and optimization described here outlines a novel coupled PHGDH assay with robust performance and improved enzymatic turnover. This protocol was developed to avoid many of the fluorescent artifacts and issues common to classic NADH-linked dehydrogenase detection methods. This novel assay enabled a large-scale ultra high-throughput screen that yielded validated small molecule PHGDH inhibitors, including one particular chemotype of interest. The systematic structural exploration and optimization of this class of 2-pyridinyl-*N*-(4-aryl)piperazine-1-carbothioamides with parallel SAR and SPR campaigns working in concert led to the discovery of NCT-503 (**31**) as a selective probe for the detailed evaluation of human PHGDH biology both *in vitro* and *in vivo.* Emerging from a structural class originally discovered and optimized for the inhibition of bacterial phosphopantetheinyl transferase (PPTase), this scaffold was selectively repurposed and optimized for PHGDH activity. Along these lines, a panel of small molecules synthesized as part of PPTase SAR campaign was tested against both enzymes, and no correlation in activity was observed between the two targets (Figure 5). In addition to an increase in potency over both the hit molecule and that of NCT-502 (**27**), moderate improvements in both solubility and microsomal stability were also observed for NCT-503 (**31**) (Table 8).

**Table 8.**
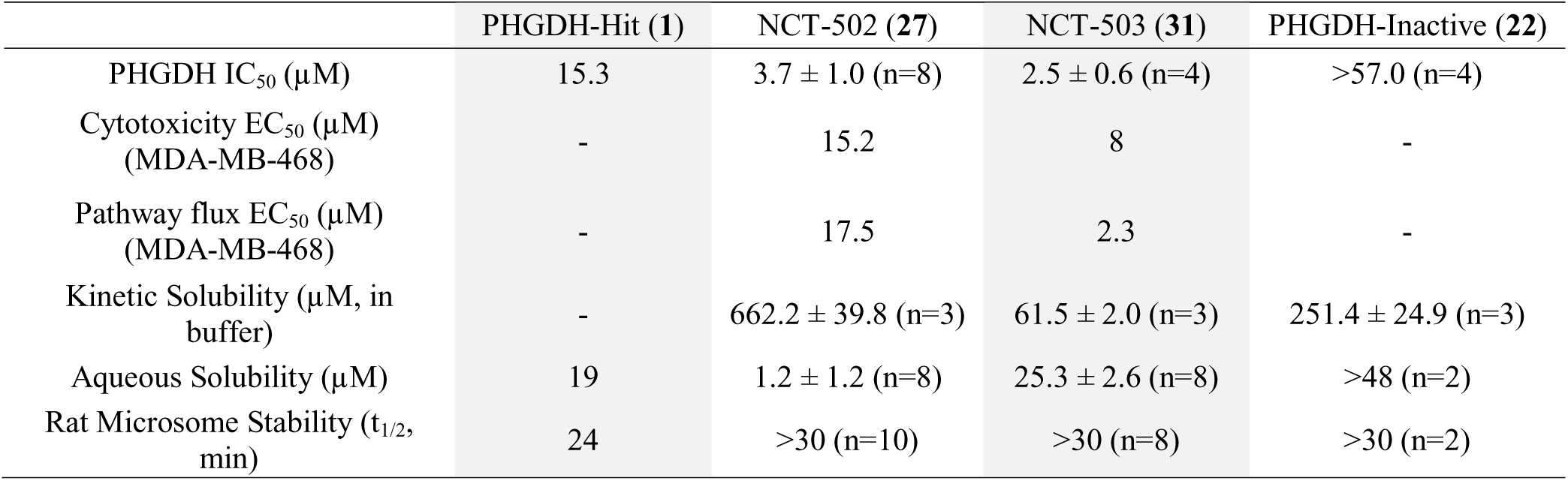
Potencies and ADME properties of PHGDH modulators

|  | PHGDH-Hit (**1**) | NCT-502 (**27**) | NCT-503 (**31**) | PHGDH-Inactive (**22**) |
| --- | --- | --- | --- | --- |
| PHGDH IC<sub>50</sub> (μM) | 15.3 | 3.7 ± 1.0 (n=8) | 2.5 ± 0.6 (n=4) | >57.0 (n=4) |
| Cytotoxicity EC<sub>50</sub> (μM)<br>(MDA-MB-468) | - | 15.2 | 8 | - |
| Pathway flux EC<sub>50</sub> (μM)<br>(MDA-MB-468) | - | 17.5 | 2.3 | - |
| Kinetic Solubility (μM, in<br>buffer) | - | 662.2 ± 39.8 (n=3) | 61.5 ± 2.0 (n=3) | 251.4 ± 24.9 (n=3) |
| Aqueous Solubility (μM) | 19 | 1.2 ± 1.2 (n=8) | 25.3 ± 2.6 (n=8) | >48 (n=2) |
| Rat Microsome Stability (t<sub>1/2</sub>,<br>min) | 24 | >30 (n=10) | >30 (n=8) | >30 (n=2) |

**Figure 5.**
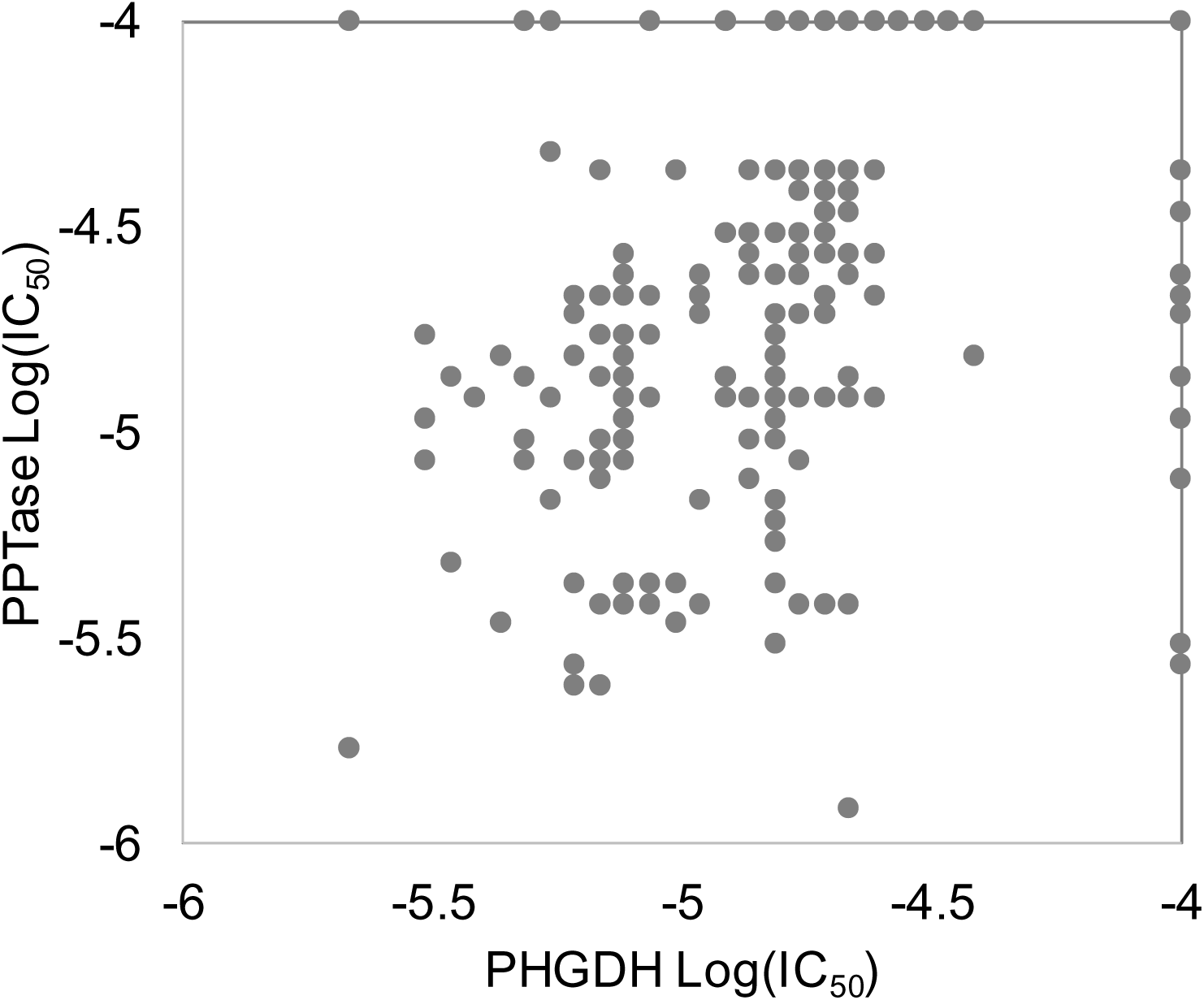
Correlation plot showing activity (IC_50_s) of a panel of analogs against both PHGDH and PPTase.

These tool compounds complement an existing set of hPHGDH probes^3^ (Figure 1) discovered through a variety of techniques including high-throughput screening^6^, fragment-based lead generation^5^, structure-based drug design^10^, as well as a merger of both fragment screening and pharmacophore convergence^11^. This class of inhibitors has expanded our understanding of the role of PHGDH in tumor metabolism.

## Experimental Procedures

### Materials

3-Phosphoglycerate (P8877), NAD^+^ (N0632), glutamate (49621), DL-glyceraldehyde 3-phosphate (G5251), glycerol-3-phosphate (G7886), TCEP (646547), *C. kluyveri* diaphorase (D5540), and resazurin (R7017) were from Sigma.

### Protein overexpression and purification

PHGDH, PSAT1 and PSPH were expressed and purified as previously described in Pacold et al, Nature Chemical Biology 12, 452–458 (2016).

cDNAs to human PHGDH, PSAT1 and PSPH were PCR amplified from human liver cDNA prepared from human liver mRNA using SuperScript III (Life Technologies) and cloned into pET30-2 with an N-terminal 6xHis Tag. Proteins were expressed in Rosetta (DE3)pLysS *E. coli* (EMD Millipore) grown to an OD of 0.6 and induced with 1 mM IPTG for 16 h at 16 °C. Bacteria were lysed at 4 °C in a French press and purified by Ni^2+^ affinity chromatography on a 5 mL HiTrap chelating HP column (GE Healthcare) attached to an AktaPURE FPLC system (GE Healthcare) using a gradient of 0–500 mM imidazole in 50 mM Na-Phosphate pH 8 and 300 mM NaCl. Peak fraction purity was assessed by SDS gel electrophoresis. Pure fractions were combined, concentrated in 15 mL UltraFree 30 concentrators (EMD Millipore) and loaded onto a HiLoad Superdex 200 prep grade 16/60 column equilibrated in 20 mM Tris pH 7.4, 100 mM NaCl, and 1 mM TCEP. Peak fractions were concentrated to [protein] ≥ 5 mg/mL, flash frozen in liquid nitrogen, and stored at −80 °C before use.

### Enzyme assays

PHGDH assay buffer contained 50 mM TEA pH 8.0, 10 mM MgCl_2_, 0.05% bovine serum albumin (BSA), and 0.01% Tween-20. PHGDH enzyme buffer consisted of assay buffer with 10 nM PHGDH and 0.15 mM NAD. PHGDH substrate buffer contained assay buffer with 10 μM EDTA, 0.625 mM glutamate, 0.05 mM 3-phosphoglycerate, 0.1 mM resazurin, 500 nM PSAT1, 500 nM PSPH and 0.1 mg/mL diaphorase. The PHGDH primary assay was performed in 1,536-well black solid-bottom plates. Substrate buffer was dispensed into each well (3 μL/well) using a BioRAPTR Flying Reagent Dispenser (Beckman Coulter), and 23 nL of DMSO-solubilized compounds were transferred into each well using a Kalypsys 1,536-well pintool equipped with 20 nL notched transfer pins. PHGDH enzyme buffer was then dispensed into each well (1 μL/well) and plates were spun in a plate centrifuge for 10 seconds to mix reagents begin the reaction. Plates were stored at room temperature and resulting fluorescence was read at 0 min 5 mg/mL, flash frozen in and 20 min using a ViewLux uHTS Microplate Imager (PerkinElmer), using a 525/20 excitation and 598/25 emission filter with a 5 sec exposure and excitation energy of 5000.

The uncoupled variation of the PHGDH assay was optimized to run in the absence of PSAT1 and PSPH downstream enzymes.

PHGDH inhibition data was analyzed by calculating delta RFUs (increase in resorufin fluorescence between *T* = 0 and *T* = 20 min) and normalizing to the delta RFU values of 0× and 1× PHGDH enzyme controls as 100% and 0% PHGDH inhibition, respectively. 32 wells of each 1536-well assay plate were dedicated to each of these 0× and 1× PHGDH controls. These normalized dose-response values were plotted in GraphPad Prism and fit using a Sigmoidal dose− response (variable slope) equation. IC_50_s for each replicate were then averaged to determine average IC_50_s with SD.

### General Synthetic Methods

All air− or moisture-sensitive reactions were performed under positive pressure of nitrogen with oven-dried glassware. Anhydrous solvents such as dichloromethane, *N,N*-dimethylformamide (DMF), acetonitrile, methanol, and triethylamine were purchased from Sigma-Aldrich. Preparative purification was performed on a Waters semi-preparative HPLC system. The column used was a Phenomenex Luna C18 (5 micron, 30 × 75 mm) at a flow rate of 45 mL/min. The mobile phase consisted of acetonitrile and water (each containing 0.1% trifluoroacetic acid). A gradient of 10% to 50% acetonitrile over 8 minutes was used during the purification. Fraction collection was triggered by UV detection (220 nM). Analytical analysis was performed on an Agilent LC/MS (Agilent Technologies, Santa Clara, CA). Purity analysis was determined using a 7 minute gradient of 4% to 100% acetonitrile (containing 0.025% trifluoroacetic acid) in water (containing 0.05% trifluoroacetic acid) with an 8 minute run time at a flow rate of 1 mL/min. A Phenomenex Luna C18 column (3 micron, 3 × 75 mm) was used at a temperature of 50 °C using an Agilent Diode Array Detector. Mass determination was performed using an Agilent 6130 mass spectrometer with electrospray ionization in the positive mode. ^1^H NMR spectra were recorded on Varian 400 MHz spectrometers. Chemical shifts are reported in ppm with nondeuterated solvent (DMSO-*h*_6_ at 2.50 ppm) as internal standard for DMSO-*d*_6_ solutions. All of the analogs tested in the biological assays have a purity greater than 95% based on LCMS analysis. High-resolution mass spectrometry was recorded on Agilent 6210 Time-of-Flight LC/MS system. Confirmation of molecular formulae was accomplished using electrospray ionization in the positive mode with the Agilent Masshunter software (version B.02).

### Experimental procedures

*N*-(4,6-dimethylpyridin-2-yl)-4-(pyridin-4-yl)piperazine-1-carbothioamide (**22**)

To a solution of di(1H-imidzazol-1-yl)methanethione (1.64 g, 9.19 mmol) in THF (24 mL) was added 4,6-dimethylpyridine-2-amine (1.12 g, 9.19 mmol). The resulting reaction mixture was heated with stirring at 40 °C for 35 min. During the reaction, the mixture was sonicated in order to produce a homogeneous yellow slurry. To the resulting mixture was added 1-(pyridin-4-yl)piperazine (1.50 g, 9.19 mmol). The resulting reaction mixture became red and was heated with stirring at 50 °C for 2 hr, then at 70 °C for 0.5 hr. The reaction mixture was diluted with water and dichloromethane. The layers were separated and the aqueous layer was re-extracted with dichloromethane. The combined organic layers were dried with MgSO_4_ and concentrated *in vacuo.* The resulting residue was taken up in DMSO and purified via reversed phase column chromatography (0 to 50% acetonitrile/water 0.1% TFA). The pure fractions were combined and most of the organic portion removed *in vacuo*. To the resulting mixture was added dichloromethane as well as saturated aqueous sodium bicarbonate solution, in order to free base the product. The layers were separated and the aqueous layer was re-extracted with dichloromethane three additional times. The combined organic layers were dried with MgSO_4_ and concentrated *in vacuo* to afford *N*-(4,6-dimethylpyridin-2-yl)-4-(pyridin-4-yl)piperazine-1-carbothioamide (**22**, 1.33 g, 44%) as a light yellow solid. ^1^H NMR (400 MHz, DMSO-*d*_6_) δ 9.71 (br s, 1H), 8.18 – 8.12 (m, 2H), 7.20 (s, 1H), 6.81 – 6.70 (m, 3H), 4.05-3.95 (m, 4H), 3.47 – 3.39 (m, 4H), 2.33 (s, 3H), 2.21 (s, 3H); ^13^C NMR (101 MHz, DMSO-*d*_6_) 181.60, 155.19 (br), 154.29, 154.03, 150.25, 148.45 (br), 119.27, 115.87, 108.47, 47.93, 45.06, 23.47, 21.07; HRMS: *m/z* (M+H)^+^ = 328.1597 (Calculated for C_17_H_22_N_5_S = 328.1590), Retention time: 1.953 min.

*N*-(4,6-dimethylpyridin-2-yl)-4-(5-(trifluoromethyl)pyridin-2-yl)piperazine-1-carbothioamide (NCT-502, **27**)

To a solution of di(1H-imidzazol-1-yl)methanethione (0.60 g, 3.4 mmol) in THF (15 mL) was added 4,6-dimethylpyridine-2-amine (0.41 g, 3.4 mmol). The resulting reaction mixture was heated with stirring at 40 °C for 30 min. During the reaction, the mixture was sonicated in order to produce a homogeneous yellow slurry. To the resulting mixture was added 1-(5-(trifluoromethyl)pyridin-2-yl)piperazine (0.78 g, 3.4 mmol). The resulting reaction mixture was heated with stirring at 50 °C for 1 hr. The reaction mixture was concentrated under a stream of air. The resulting residue was taken up in DMSO and purified via reverse phase column chromatography (acetonitrile/water 0.1% HCl). Combined fractions were partially concentrated *in vacuo*, neutralized with saturated aqueous sodium bicarbonate solution, and filtered to remove the solid. The solid was taken up in DMSO and repurified via reverse phase column chromatography (5 to 100% acetonitrile/water 0.1% TFA). Combined fractions were partially concentrated *in vacuo* (to remove organics) and filtered to provide the desired product, *N*-(4,6-dimethylpyridin-2-yl)-4-(5-(trifluoromethyl)pyridin-2-yl)piperazine-1-carbothioamide trifluoroacetate salt (NCT-502, **27**), 0.44 g, 26 %), as a solid. ^1^H NMR (400 MHz, DMSO-*d*_6_) δ 10.02 (br s, 1H), 8.42 (m, 1H), 7.82 (dd, *J* = 9.2, 2.6 Hz, 1H), 7.27 (s, 1H), 6.97 – 6.90 (m, 2H), 4.06 – 3.99 (m, 4H), 3.80 – 3.72 (m, 4H), 2.41 (s, 3H), 2.30 (s, 3H); ^13^C NMR (101 MHz, DMSO-*d*_6_) δ 181.34, 160.18, 158.81, 158.46, 145.68 (q, *J*_C-F_ = 5.05 Hz), 135.06 (q, *J*_C-F_ = 3.03 Hz), 126.65, 123.97, 120.6 (br), 116.93, 113.91 (q, *J*_C-F_ = 32.32 Hz), 106.72, 48.21, 43.81, 22.05, 21.37; HRMS: *m/z* (M+H)^+^ = 396.1480 (Calculated for C_18_H_21_F_3_N_5_S = 396.1464), retention time: 3.004 min.

*N*-(4,6-dimethylpyridin-2-yl)-4-(4-(trifluoromethyl)benzyl)piperazine-1-carbothioamide (NCT-503, **31**)

To a solution of *tert*-butyl piperazine-1-carboxylate (1.31 g, 7.05 mmol) and triethylamine (1.2 mL, 8.5 mmol) in THF (30 ml) was added 1-(bromomethyl)-4-(trifluoromethyl)benzene (1.69 g, 7.05 mmol). The reaction mixture was heated with stirring at 55 °C overnight. The reaction mixture was diluted with water and dichloromethane. The layers were separated and the aqueous layer was re-extracted with dichloromethane. The combined organic layers were dried with MgSO_4_ and concentrated *in vacuo* to afford *tert*-butyl 4-(4-(trifluoromethyl)benzyl)piperazine-1-carboxylate as a yellow oil; ^1^H NMR (400 MHz, DMSO-*d*_6_) δ 7.70 – 7.62 (m, 2H), 7.55 – 7.47 (m, 2H), 3.55 (s, 2H), 3.33 – 3.25 (m, 4H), 2.33 – 2.26 (m, 4H), 1.36 (s, 9H); ^13^C NMR (101 MHz, DMSO-*d*_6_) reaction was determined to 4.04 Hz), δ 79.20, 61.62, 55.33, 52.79, 28.49. This material was taken up in dichloromethane (20 mL) and treated with TFA (2 mL). After standing at rt for 1 hr, an additional aliquot of TFA (3 mL) was added. Upon standing at rt for an additional 2 hr, the reaction was determined to be complete by LCMS (LCMS: *m/z* (M+H)^+^ = 245.1). The reaction mixture was concentrated *in vacuo*, rediluted with ∼50 mL of dichloromethane, and reconcentrated *in vacuo* to yield a partially crystalline, faint tan solid which was used without further purification; ^1^H NMR (600 MHz, DMSO-*d*_6_) δ 8.84 (br s, 2H), 7.76 (d, *J* = 8.0 Hz, 2H), 7.62 (d, *J* = 7.8 Hz, 2H), 3.99 (br s, 2H), 3.19 (br s, 4H), 3.01 – 2.71 (br m, 4H); ^13^C NMR (151 MHz, DMSO-*d*_6_) δ 161.45 (q, *J* = 34.7 Hz), 133.83 (m), 128.34 (q, *J* = 49.8 Hz), 127.27 (q, *J* = 271.8 Hz), 119.09 (q, *J* = 292.9 Hz), 62.64 (m), 51.59, 44.99 (m).

To a solution of di(1H-imidzazol-1-yl)methanethione (1.26 g, 7.05 mmol) in THF (30 mL) was added 4,6-dimethylpyridine-2-amine (0.861 g, 7.05 mmol). The resulting reaction mixture was heated at 40 °C for 35 min. During the reaction, the mixture was sonicated in order to produce a homogeneous yellow slurry. This slurry was transferred to another vial containing a slurry comprised of the TFA salt of 1-(4-(trifluoromethyl)benzyl)piperazine (from step 2), THF (10 mL), and triethylamine (1 mL, 7.05 mmol). An additional aliquot of THF (5 mL) was used to complete the transfer. The resulting reaction mixture was heated with stirring at 70 °C for 1.25 hr. The reaction mixture was diluted with water and dichloromethane. The layers were separated and the aqueous layer was re-extracted with dichloromethane. The combined organic layers were dried with MgSO_4_ and concentrated *in vacuo.* The resulting residue was taken up in DMSO and purified via reversed phase column chromatography (10 to 50% acetonitrile/water 0.1% TFA). The pure fractions were combined and most of the organic portion removed *in vacuo*. To the resulting mixture was added dichloromethane as well as saturated aqueous sodium bicarbonate solution, in order to free base the product. The layers were separated and the aqueous layer was re-extracted with dichloromethane three additional times. The combined organic layers were dried with MgSO_4_ and concentrated *in vacuo* to afford *N*-(4,6-dimethylpyridin-2-yl)-4-(4-(trifluoromethyl)benzyl)piperazine-1-carbothioamide (NCT-503, **31**, 0.809 g, 28% over 3 steps) as a colorless foam. ^1^H NMR (400 MHz, DMSO-*d*_6_) δ 9.59 (br s, 1H), δ 7.72 – 7.64 (m, 2H), 7.58 – 7.48 (m, 2H), 7.13 (s, 1H), 6.70 (s, 1H), 3.91-3.79 (m, 4H), 3.60 (s, 2H), 2.44-2.38 (m, 4H), 2.32 (s, 3H), 2.20 (s, 3H); ^13^C NMR (101 MHz, DMSO-*d*_6_) δ 181.44, 154.04, 143.35 (m), 129.92, 128.85, 128.13 (q, *J*_C-F_ = 31.31 Hz), 126.15, 125.55 (q, *J*_C-F_ = 4.04 Hz), 123.44, 119.17 (br), 115.62, 61.26, 52.67, 48.88, 23.56 (br), 21.03; HRMS: *m/z* (M+H)^+^ = 409.1673 (Calculated for C_20_H_24_F_3_N_4_S = 409.1668), Retention time: 2.476 min.

## Acknowledgements

We thank Heather Baker, Danielle Bougie, Yuhong Fang, Elizabeth Fernandez, Misha Itkin, Zina Itkin, Christopher LeClair, William Leister, Crystal McKnight, and Paul Shinn for assistance with chemical purification and compound management. This research was supported by the Molecular Libraries Initiative of the National Institutes of Health Roadmap for Medical Research Grant U54MH084681 and the Intramural Research Program of the National Center for Advancing Translational Sciences, National Institutes of Health. D.M.S. is supported by the NIH (R01 CA129105, R01 CA103866, and R37 AI047389) and is an investigator of the Howard Hughes Medical Institute. M.E.P. is supported by the NIH (K22 CA212059), the Mary Kay Foundation (017-32), the V foundation (V2017-004), and the Shifrin-Myers Breast Cancer Discovery Fund at NYU.

---

**Scheme 1.**
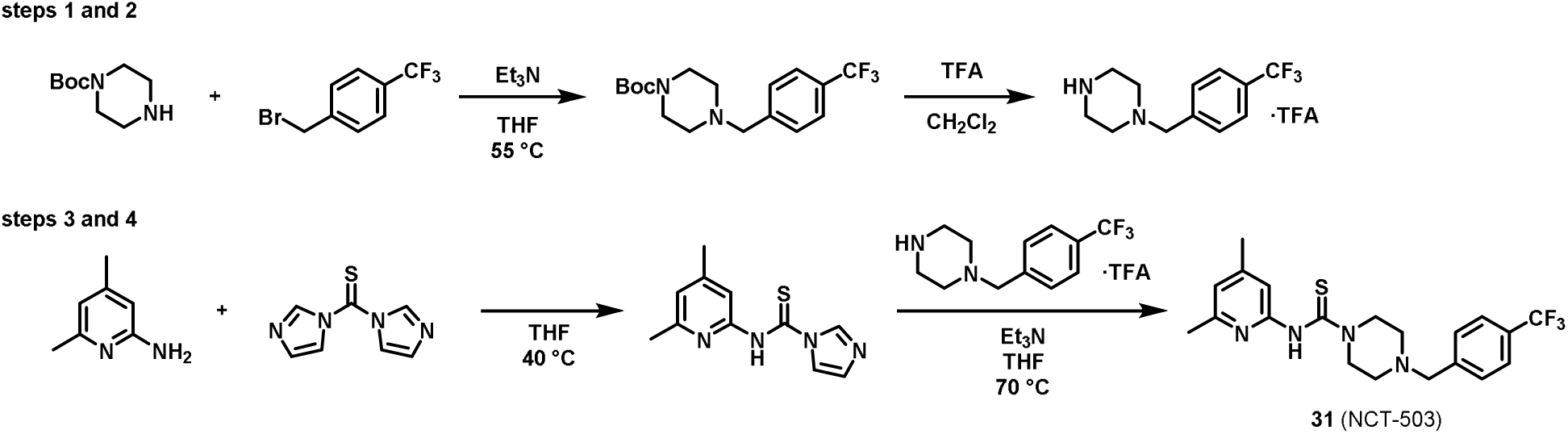
Methodology used to synthesize benzyl-substituted piperazines

